# Persistent biofluid small molecule alterations induced by *Trypanosoma cruzi* infection are not restored by antiparasitic treatment

**DOI:** 10.1101/2023.06.03.543565

**Authors:** Danya A. Dean, Jarrod Roach, Rebecca Ulrich vonBargen, Yi Xiong, Shelley S. Kane, London Klechka, Kate Wheeler, Michael Jimenez Sandoval, Mahbobeh Lesani, Ekram Hossain, Mitchelle Katemauswa, Miranda Schaefer, Morgan Harris, Sayre Barron, Zongyuan Liu, Chongle Pan, Laura-Isobel McCall

**Affiliations:** Department of Chemistry and Biochemistry, University of Oklahoma, Norman, OK, 73019, USA; Laboratories of Molecular Anthropology and Microbiome Research, University of Oklahoma, Norman, OK, 73019; USA; Department of Biomedical Engineering, University of Oklahoma, Norman, OK, 73019, USA; Department of Microbiology and Plant Biology, University of Oklahoma, Norman, OK, 73019, USA; Department of Biology, University of Oklahoma, Norman, OK, 73019, USA; Department of Neuroscience, Amherst College, Hampshire County, MA, 01002; USA

**Keywords:** Chagas disease, *Trypanosoma cruzi*, Biomarkers, Metabolites, Clinical treatment failure, Urine

## Abstract

**Table of contents graphic:** 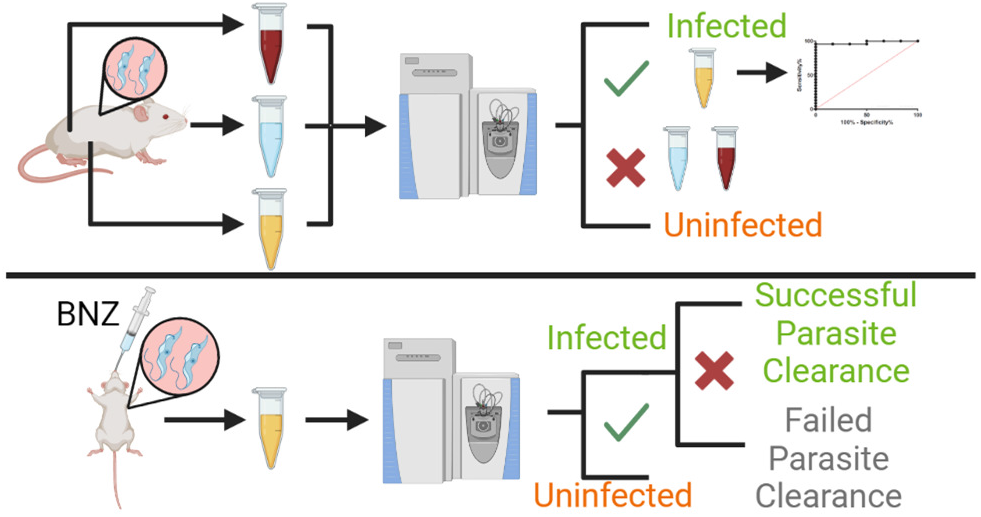

Chagas Disease (CD), caused by *Trypanosoma cruzi (T. cruzi)* protozoa, is a complicated parasitic illness with inadequate medical measures for diagnosing infection and monitoring treatment success. To address this gap, we analyzed changes in the metabolome of *T. cruzi-*infected mice via liquid chromatography tandem mass spectrometry analysis of clinically-accessible biofluids: saliva, urine, and plasma. Urine was the most indicative of infection status, across mouse and parasite genotypes. Metabolites perturbed by infection in the urine include kynurenate, acylcarnitines, and threonylcarbamoyladenosine. Based on these results, we sought to implement urine as a tool for assessment of CD treatment success. Strikingly, it was found that mice with parasite clearance following benznidazole antiparasitic treatment had comparable overall urine metabolome to mice that failed to clear parasites. These results match with clinical trial data in which benznidazole treatment did not improve patient outcomes in late-stage disease. Overall, this study provides insights into new small molecule-based CD diagnostic methods and a new approach to assess functional treatment response.

## Introduction

With 6-7 million people infected worldwide and at least 300,000 in the United States, Chagas disease (CD) is progressively becoming a worldwide concern due to migration of infected individuals ^1^. This parasitic disease is caused by the protozoan *Trypanosoma cruzi (T. cruzi),* transmitted by triatomine insects. It can also be acquired through food and drink, organ transplants, blood transfusions or congenital transmission. The disease presents in three stages: acute, indeterminate, and chronic. The acute stage presents with non-specific symptoms such as nausea and fever; however, patients become asymptomatic within a few weeks. This is the indeterminate stage, which may last decades. 30-40% of infected people progress to symptomatic chronic stage with severe cardiac symptoms such as arrhythmias, cardiac failure, and apical aneurysms, and less commonly intestinal tract symptoms including megacolon and megaesophagus ^1, 2^. 55-65% of symptomatic patients die due to cardiac arrest, 25–30% due to heart failure and 10-15% due to a thromboembolic event ^2^.

Current chronic CD diagnostic methods include several serological tests: enzyme-linked immunosorbent assays, indirect hemagglutination and indirect immunofluorescence assays ^3, 4^. These tests rely on the presence of anti-*T. cruzi* antibodies for diagnosis, but suffer from low reliability due to the rate of false positives (0.3 – 3.2%) and false negatives (0.7 – 3.7%). Indeed, there must be a minimum of two positive test results to conclusively diagnose CD. Approximately 50% of *T. cruzi-*positive blood was misdiagnosed in Venezuelan blood banks due to the low specificity and sensitivity of current diagnostic methods ^5, 6^. This is concerning, as misdiagnosed blood samples can be used for blood transfusions, ultimately leading to more *T. cruzi* infections. Additionally, microscopy methods are available but are only useful in the acute stage of CD.

Current treatments for CD include the antiparasitics benznidazole and nifurtimox. Unfortunately, these drugs have severe adverse effects, sometimes leading to treatment interruption. In addition, up to 20% of patients do not achieve full parasite clearance by these standard treatments ^7–11^. Addressing treatment success is thus essential to prioritize follow-up treatment in these patient cohorts and to guide drug development. However, traditional serological tests can take decades to become negative after parasite clearance. As a faster alternative, many clinical trials are relying on PCR techniques ^7, 12, 13^. PCR detection is unfortunately only 60-70% sensitive, as it is not consistently positive in infected individuals. PCR thus cannot indicate treatment success but only parasitological treatment failure ^14–16^. In a survey of 155 CD experts around the world, 62 acknowledged early assessment of treatment response as a high need in research priorities in the CD community ^14^.

Importantly, all these diagnostic methods also lack the ability to monitor disease progression. In the BENEFIT clinical trial, while successful parasitological cure was achieved with benznidazole treatment, benznidazole-treated patients nevertheless went into cardiac failure or died at comparable rates to placebo controls, indicating that parasite burden alone cannot predict clinical outcomes ^12^. While electrocardiograms may be useful for identifying heart dysfunction, they cannot diagnose chronic stage CD but simply cardiac disease ^17^. Furthermore, most patients in the chronic stage of CD have irreversible cardiac fibrosis, which highlights the importance of earlier diagnosis. Brain Natriuretic Peptide (BNP), a known biomarker of heart failure, is increased in patients with CD. It identifies cardiac disease but does not necessarily diagnose parasitic infection or predict progression to severe CD from the indeterminate stage ^18, 19^. Researchers have also tested magnetic resonance imaging (MRI) on CD patients to assess disease severity, but this was mainly effective in already-symptomatic patients ^20^.

There is therefore a strong need for novel diagnostic methods for CD. Metabolomics can bridge this gap. Metabolomics is the integration of analytical and biochemical methods to study small molecules, broadly defined here as any molecule with a size of 1500 Da or less. These molecules are the products of metabolic pathways, critical signaling molecules, and the building blocks of biology, including amino acids, nucleotides and lipids. As such, they are direct markers of changes in biochemical processes ^21–23^. We hypothesized that liquid chromatography tandem mass spectrometry (LC-MS/MS)-based metabolomics of biofluids will enable the discovery of CD biomarkers that can monitor chronic *T. cruzi* infection status, disease progression and treatment success. We found that changes in the urine metabolome were most reflective of disease progression, with common biomarkers validated between replicate infection cohorts and in two different mouse strain and parasite combinations, across two *T. cruzi* discrete typing units (DTUs) ^24^. Specific perturbed metabolites include kynurenate, acylcarnitines, and threonylcarbamoyladenosine. Strikingly, the urinary metabolome did not re-normalize following benznidazole treatment, matching with clinical conclusions from the BENEFIT trial ^12^. Overall, these results suggest that clinical treatment failure may be associated with an inability to restore metabolism post-treatment, and that urine metabolomics may be a faster way to monitor disease progression and the functional differences between treatment success vs treatment failure.

## Results

### *T. cruzi* infection perturbs the urine, saliva and plasma metabolome

Building on our prior work demonstrating that the cardiac metabolome differed between mild and severe *T. cruzi i*nfection ^25^, we assessed whether metabolome alterations were also observed in clinically-accessible biofluids (saliva, urine and plasma) over the course of experimental *T. cruzi* infection with parasite strain Sylvio X10/4 in male Swiss Webster mice. Control samples were also collected from mice treated with isoproterenol to induce cardiac hypertrophy independent of *T. cruzi* ^26^. We confirmed that isoproterenol treatment induced persistent heart damage in this model via qRT-PCR assessment of *Bnp* gene expression (average 2^-ΔΔCt^=2.71±0.62 compared to *Gapdh* expression and to uninfected samples at 90 day timepoint; p=0.029, Student’s T test, compared to uninfected samples for ΔCt values). As expected, cardiac parasite burden was highest in the acute stage of infection and then decreased to low levels (**Fig. 1A**). *Bnp* gene expression levels, in contrast, were only significantly affected at the chronic (90 days post-infection (DPI)) timepoint in infected animals compared to uninfected animals (**Table 1**).

**Fig. 1.**
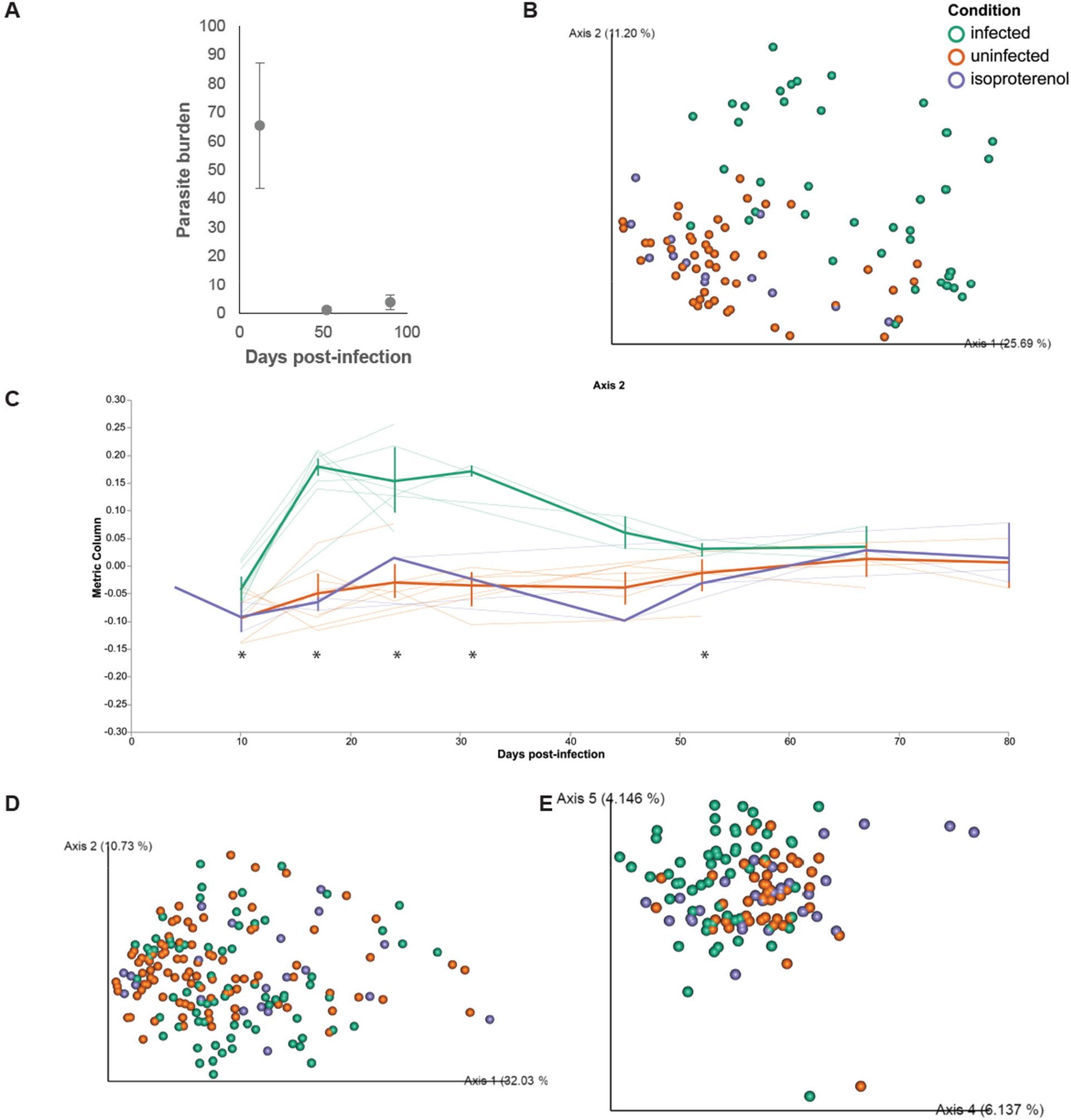
*T. cruzi* infection impacts the urine metabolome more than the salivary and plasma metabolome. (**A**) Parasite burden at heart base decreases as mice transition from acute to chronic infection. (**B**) Urine sample principal coordinate analysis. (**C**) Urine sample volatility analysis showing separation between infected samples and both uninfected and isoproterenol-treated samples. Thick lines indicate group mean and thin lines represent the trajectory of each individual mouse along principal coordinate axis 2. *, PERMANOVA p<0.05 for infected to uninfected animals. (**D**) Saliva sample principal coordinate analysis. (**E**) Plasma sample principal coordinate analysis.

**Table 1.** Impact of infection on cardiac *Bnp* expression.

| Timepoint | $2^{-\Delta\Delta Ct} \pm$ standard error (normalized to *Gapdh* and to uninfected samples) | Student's T test p-value, two-tailed, for infected $\Delta Ct$ to uninfected group $\Delta Ct$ at matched timepoints |
| --- | --- | --- |
| 12 DPI | $2 \pm 0.92$ | 0.44 |
| 52 DPI | $2.98 \pm 1.08$ | 0.19 |
| 90 DPI | $7.59 \pm 4.5$ | 0.047 |

The overall urine metabolome was the most impacted by infection, followed by saliva and plasma (**Fig. 1B-E** and **Figure S1**, PERMANOVA p<0.05 R^2^=0.22 at day 28 for plasma, all other timepoints non-significant; R^2^ range for significant timepoints for saliva from R^2^=0.13 at day 25 to 0.18 at day 18; R^2^ range for significant timepoints for urine from 0.26 at day 10 to 0.44 at day 52). Strikingly, the overall impact of infection on the saliva metabolome was similar to the impact of isoproterenol treatment (**Fig. S1A**; both groups, PERMANOVA p<0.05 to uninfected untreated at day 11 and day 18), indicating comparable effects on the metabolome. In contrast, the urine metabolome of infected mice diverged from both isoproterenol-treated and uninfected animals, indicating effects specific to *T. cruzi* infection (**Fig. 1BC**).

Next, we assessed the specific metabolites perturbed in each biofluid over time using generalized linear mixed models (GLMM). Based on GLMM analysis, urine revealed the most metabolites perturbed by infection (182 metabolites), followed by saliva (37 metabolites), then plasma (17 metabolites) (**Table 2, Table S1**). However, there were very few metabolites perturbed by isoproterenol treatment (**Table 2**). We then sought to determine commonalities between biofluids and between infected and isoproterenol samples for a given biofluid (**Fig. 2**). Interestingly, there was limited overlap between biofluids for metabolites perturbed by *T. cruzi* infection or by isoproterenol treatment (**Fig. 2AB**). In addition, there was no commonality found in plasma and in urine between metabolites perturbed by infection and by isoproterenol, and only 1 metabolite overlapped in saliva samples (**Fig. 2CDE**). The one metabolite commonly perturbed by both interventions in saliva samples was an analog of omega-hydroxydodecanoate (*m/z* 244.1904, retention time (RT) 4.84 min). The common metabolite perturbed by infection in both saliva and urine was acetylcarnitine (*m/z* 204.123, RT 0.39 min) (**Table 2**). Metabolites specifically impacted in the saliva include phenylalanine (*m/z* 166.0863 RT 0.63 min, increased by infection), carnitine (*m/z* 162.1124 RT 0.3 min) and acetylcarnitine (*m/z* 204.123 RT 0.39 min, **Table S1, Fig. S2**). In the plasma, annotated infection-impacted metabolites include kynurenine (*m/z* 209.092 RT 0.65 min, increased by infection) and N-Acetyl-L-Leucine (*m/z* 174.1125 RT 2.77 min). In contrast, palmitoylcarnitine (*m/z* 400.3417 RT 6.6 min) was impacted by isoproterenol treatment (**Table S1, Fig. S3**).

**Fig. 2.**
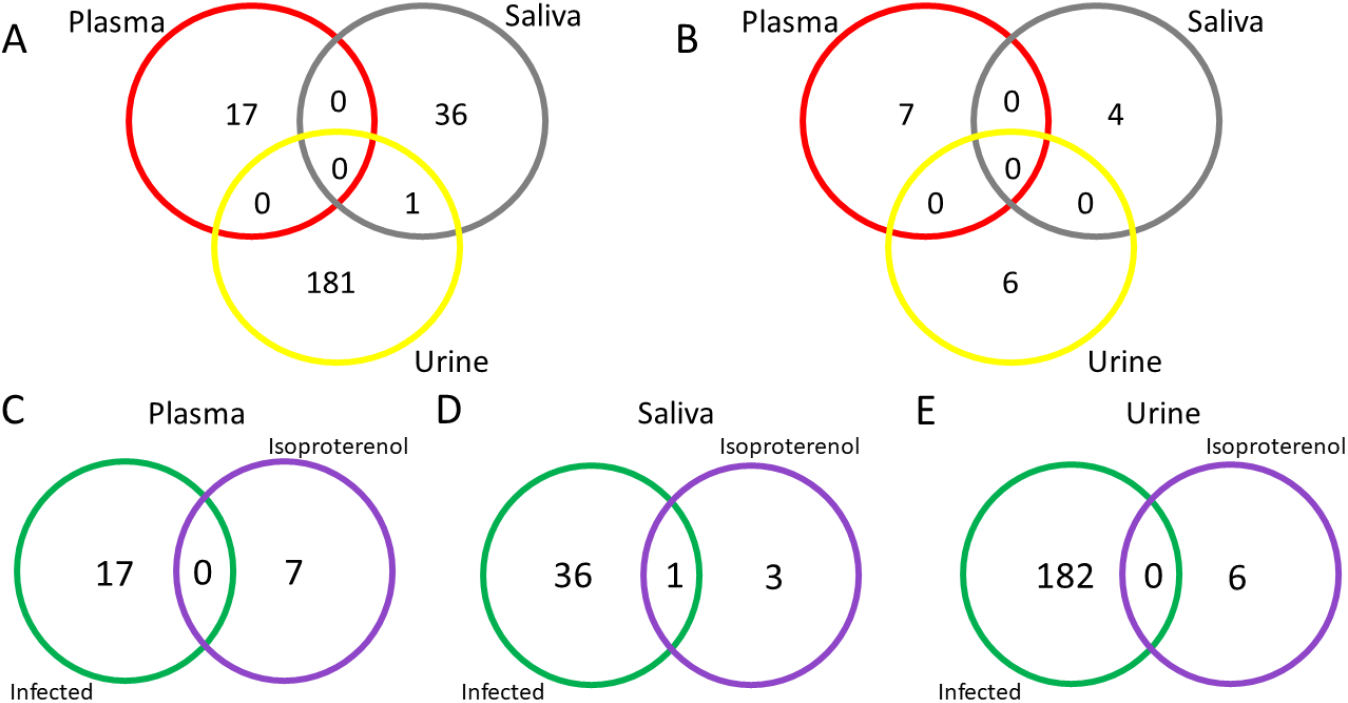
Lack of commonalities between biofluids or in a given biofluid in response to isoproterenol treatment or *T. cruzi* infection, based on GLMM output. (**A**) Limited overlap of metabolites perturbed by *T. cruzi* infection between biofluids. (**B**) No overlap of metabolites perturbed by isoproterenol treatment between biofluids. (**C**) No commonalities between metabolites significantly perturbed by infection or isoproterenol treatment in plasma samples. (**D**) Limited commonality between significantly perturbed metabolites in saliva samples. (**E**) No commonalities between significantly perturbed metabolites in urine samples.

**Table 2.**
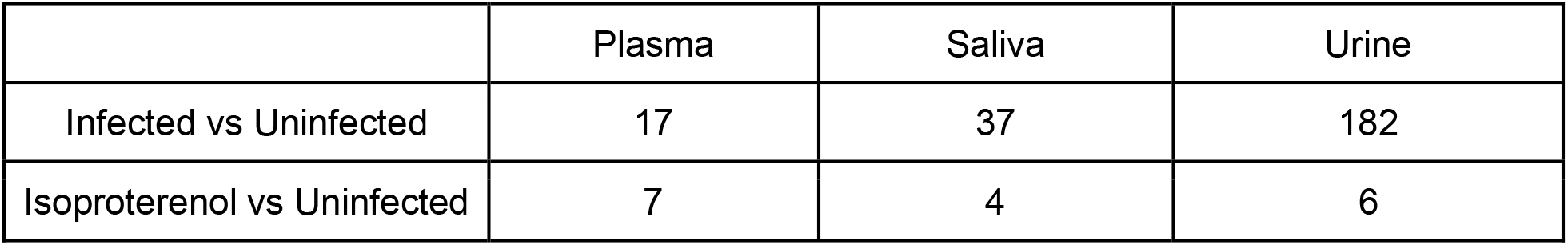
Number of significant metabolites perturbed by infection or by isoproterenol treatment in each biofluid over time.

|  | Plasma | Saliva | Urine |
| --- | --- | --- | --- |
| Infected vs Uninfected | 17 | 37 | 182 |
| Isoproterenol vs Uninfected | 7 | 4 | 6 |

To determine whether the limited commonality of metabolites identified was due to inherent heterogeneity between sample types, we developed Venn diagrams of all detected metabolites, irrespective of abundance (**Fig. 3**). Large commonality was found between biofluids in both infected and isoproterenol-treated samples in terms of metabolite presence vs absence, with greater overlap between plasma and urine than between plasma and saliva (Fisher’s exact test p<0.05; **Fig. 3AB**). The same large overlap was also noticed in each individual biofluid between interventions (**Fig. 3C-E**). This indicates that metabolites are mostly present across biofluids, but significant differences in response to perturbations are observed between biofluids and between interventions.

**Fig. 3.**
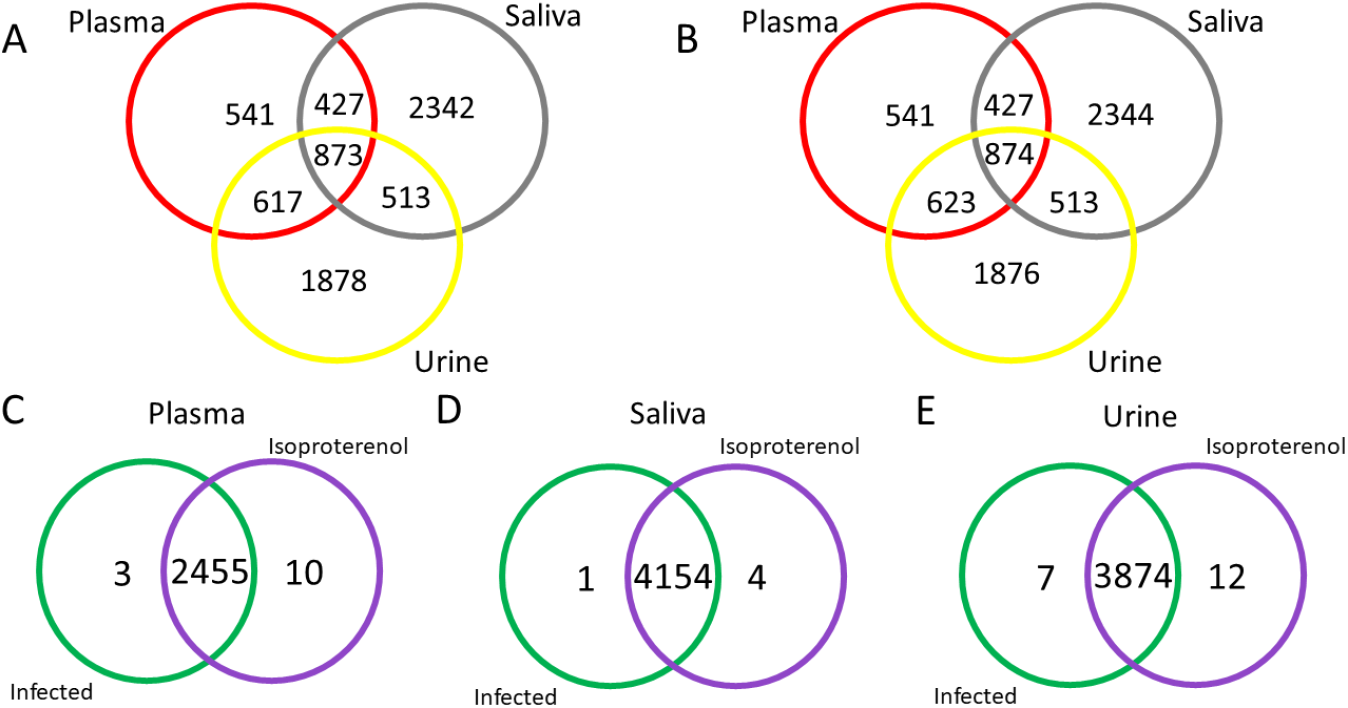
Strong commonalities between biofluids in terms of metabolite occurrence. (**A**) Large overlap of metabolites present in infected biofluids. (**B**) Large overlap of metabolites present following isoproterenol injection in biofluids. (**C**) Large commonalities found between metabolites in infected or isoproterenol treated plasma samples. (**D**) Large commonalities between metabolites in infected or isoproterenol treated saliva samples. (**E**) Large commonalities between metabolites in infected or isoproterenol treated urine samples.

Given that urine showed the strongest response to infection, we focused subsequent analysis on this biofluid. First, we validated by targeted MS analysis a subset of the metabolites output by GLMM in our initial untargeted analysis, on an independent infection cohort of male and female Swiss Webster mice infected with *T. cruzi* strain Sylvio X10/4 (same mouse and parasite strain as discovery cohort). None of these features overlapped with those perturbed by isoproterenol treatment (**Table S1**). One-third of the significant features from the discovery cohort also showed statistical significance (FDR-corrected Mann-Whitney p<0.05) and the same direction of change to uninfected samples in the validation cohort, with slightly better validation in males compared to females (17% in females and 33% in males, **Table 3**). This is likely due to the fact that our discovery cohort was male mice. However, *m/z* 272.1488 RT 3.77 min, *m/z* 298.2007 RT 3.94 min and *m/z* 311.069 RT 3.26 min were already significantly different at the pre-infection timepoints in females and thus would not be suitable biomarkers. Only a minority were significant but with the opposite direction of fold change compared to the discovery cohort (11% in females and 5% in males). One-third of the features from the discovery cohort were not observed in the validation cohort (24% in females and 30% in males), with the remainder of the features detected but showing no significant difference between infected and uninfected samples (28/63 in females and 20/63 in males). Such reproducibility rates are comparable to other urine metabolomics studies (e.g. ^27, 28^).

**Table 3.** Validated biomarkers.

| *m/z* | RT (min) | Timepoints with $p \leq 0.05$ (Mann-Whitney, FDR-corrected) and matching discovery cohort, in females<sup>1,2</sup> | Timepoints with $p \leq 0.05$ (Mann-Whitney, FDR-corrected) and matching discovery cohort, in males<sup>1,2</sup> | Annotation<sup>3</sup> |
| --- | --- | --- | --- | --- |
| 134.06 | 1.83 | Not detected | Week 10 (opposite pattern at 1 week) | Indoxyl-containing molecule (4 ppm) |
| 151.144 | 0.3 | Week 17 | Week 2, 4, 5, 10, 18 | NA |
| 176.0916 | 0.51 | Week 2, 3 | Week 2, 4, 5 | NA |
| 214.1071 | 2.55 | Week 2 | Week 1, 2, 5, 18 | NA |
| 220.0636 | 3.49 | Not consistently detected past day 0 | Week 1, 2, 4, 5 | NA |
| 224.1279 | 2.82 | N/S | Week 1, 4, 5 | NA |
| 242.1383 | 3.5 | N/S | Week 4, 5 | NA |
| 247.1073 | 3.5 | Week 2, 3, 6 | N/S | NA |
| 264.0377 | 0.38 | N/S | Week 5 | NA |
| 272.1488 | 3.77 | Not suitable as a biomarker: already significant at week 0 (also significant at week 6, 10) | Not detected | NA |
| 280.165 | 0.3 | Week 2, 5, 6, 10, 17 | Week 1, 2, 4, 5, 10, 18 | NA |
| 298.2007 | 3.94 | Not suitable as a biomarker: already significant at week 0. | Week 2, 5 | NA |
| 307.201 | 4.57 | N/S | Week 1, 10, 18 | NA |
| 311.069 | 3.26 | Not suitable as a biomarker: already significant at week 0 (also significant at week 1, 2, 10, 17) | Week 5 | NA |
| 333.0509 | 3.26 | N/S | Week 1, 5 | NA |
| 343.222 | 3.64 | Week 5 | Week 4, 5 | NA |
| 375.114 | 0.52 | Week 3 | Week 1, 2, 4, 10 | NA |
| 382.258 | 4.94 | N/S | Week 4, 10 | CAR 14:3;O (3 ppm) |
| 400.2687 | 4.35 | N/S | Week 10 | NA |
| 411.1543 | 3.53 | Week 5, 10, 17 | Week 10 | NA |
| 413.141 | 2.37 | Week 1, 2, 3, 5 | Week 10 | N6-Threonylcarbamoyl adenosine (3 ppm) |
| 434.2378 | 4.35 | N/S | Week 10 | NA |
| 497.1334 | 3.1 | Week 3 (opposite pattern) | Week 2, 5 | NA |

<sup>1</sup> N/S, non-significant.
<sup>2</sup> Female samples were collected on weeks 0, 1, 2, 3, 5, 6, 10, and 17, while male samples were collected on weeks 0, 1, 2, 4, 5, 10, and 18, for logistical reasons.
<sup>3</sup> All annotations at level 2/3 confidence <sup>29</sup>.

For the features that showed similar patterns in both males and females, we performed ROC (Receiver operating characteristic) analysis at the matching timepoints. Overall, we observed excellent Area Under the Curve (AUC) values, with a range of 0.8318 to 1 (**Fig. 4**).

**Fig. 4.**
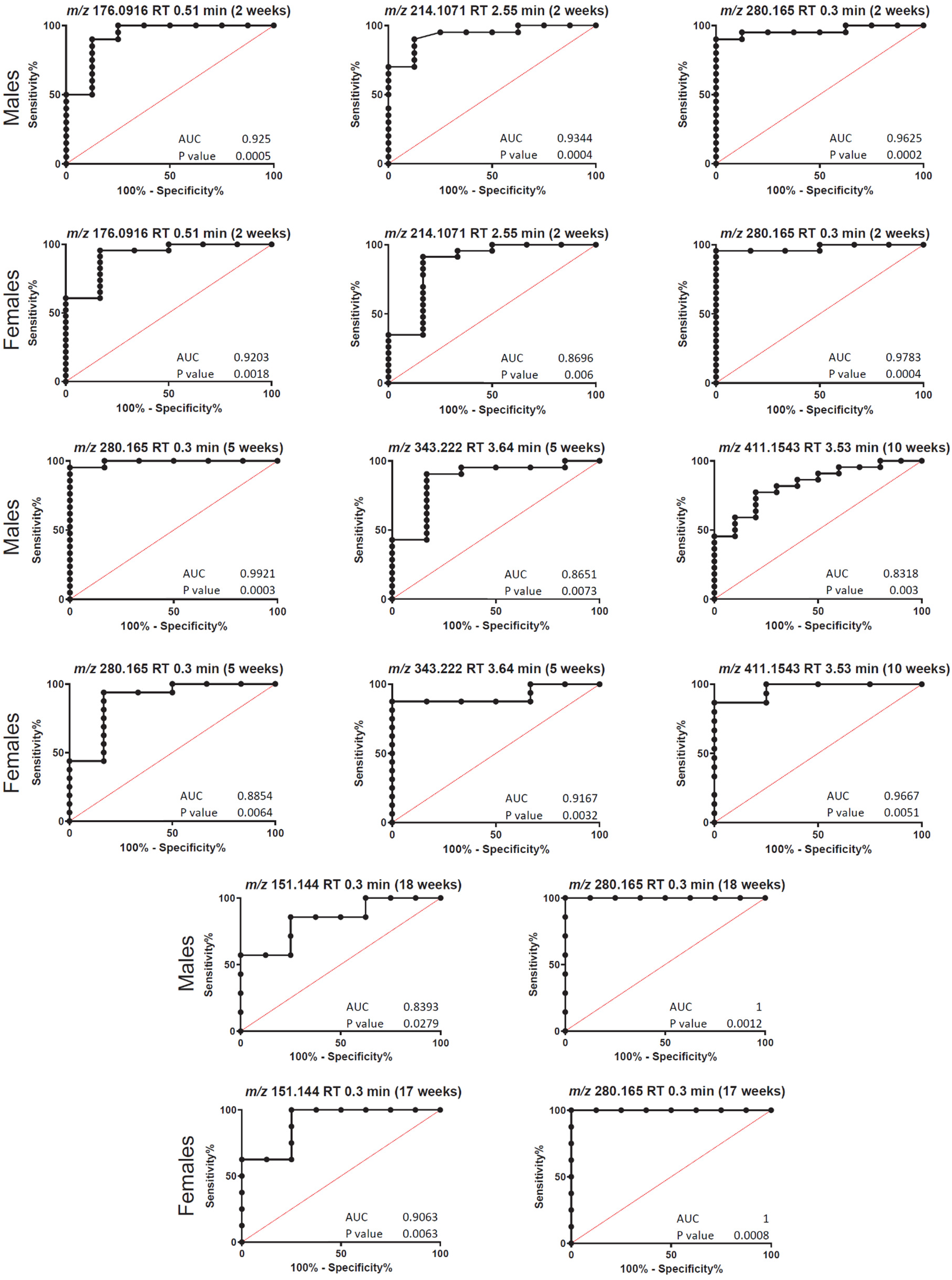
Representative ROC curves (validation cohort).

### Infection-induced alterations in the urinary metabolome are not restored by antiparasitic treatment

Having established that the urinary metabolome is perturbed by infection, we then assessed whether this could be restored by parasite clearance. To enable this analysis, we switched to a well-characterized model of antiparasitic treatment assessment, female BALB/c mouse infection with luciferase-expressing strain CL Brener ^30, 31^. For comparison of parasite clearance vs parasite persistence, a parasite burden cutoff post-immunosuppression of uninfected group value + 3 standard deviations was used (20 parasite equivalents). By this definition, 17 mice were deemed to have successfully cleared parasites, and 25 were deemed parasitological treatment failures (**Fig. 5A**). There was a clear and significant difference in the overall urine metabolome between the uninfected controls and the infected mice by principal coordinate analysis (whether treatment cleared parasites or not, PERMANOVA p<0.05, R^2^ = 0.08, **Fig. 5B**), confirming our prior findings in a second mouse strain-parasite strain combination, in a divergent DTU ^24^ (compare **Fig. 1B** to **Fig. 5B**). Note that these samples were collected prior to immunosuppression, to avoid confounding effects from the cyclophosphamide treatment. In contrast, there was no significant difference between the mice that successfully cleared parasites and mice that failed to do so, at 35 days post-treatment (PERMANOVA p>0.05, **Fig. 5C**). Likewise, no proportional correlation to post-immunosuppression parasite burden was observed (**Fig. 5D**). Fifteen of the features in **Table 3** were detected in this independent infection system, four of which significantly differed between infected and uninfected samples but did not show any restoration even following sterile cure by benznidazole (Kruskal Wallis p<0.05 with post-hoc Dunn’s test, FDR-corrected, **Fig. 6A-D**).

**Fig. 5.**
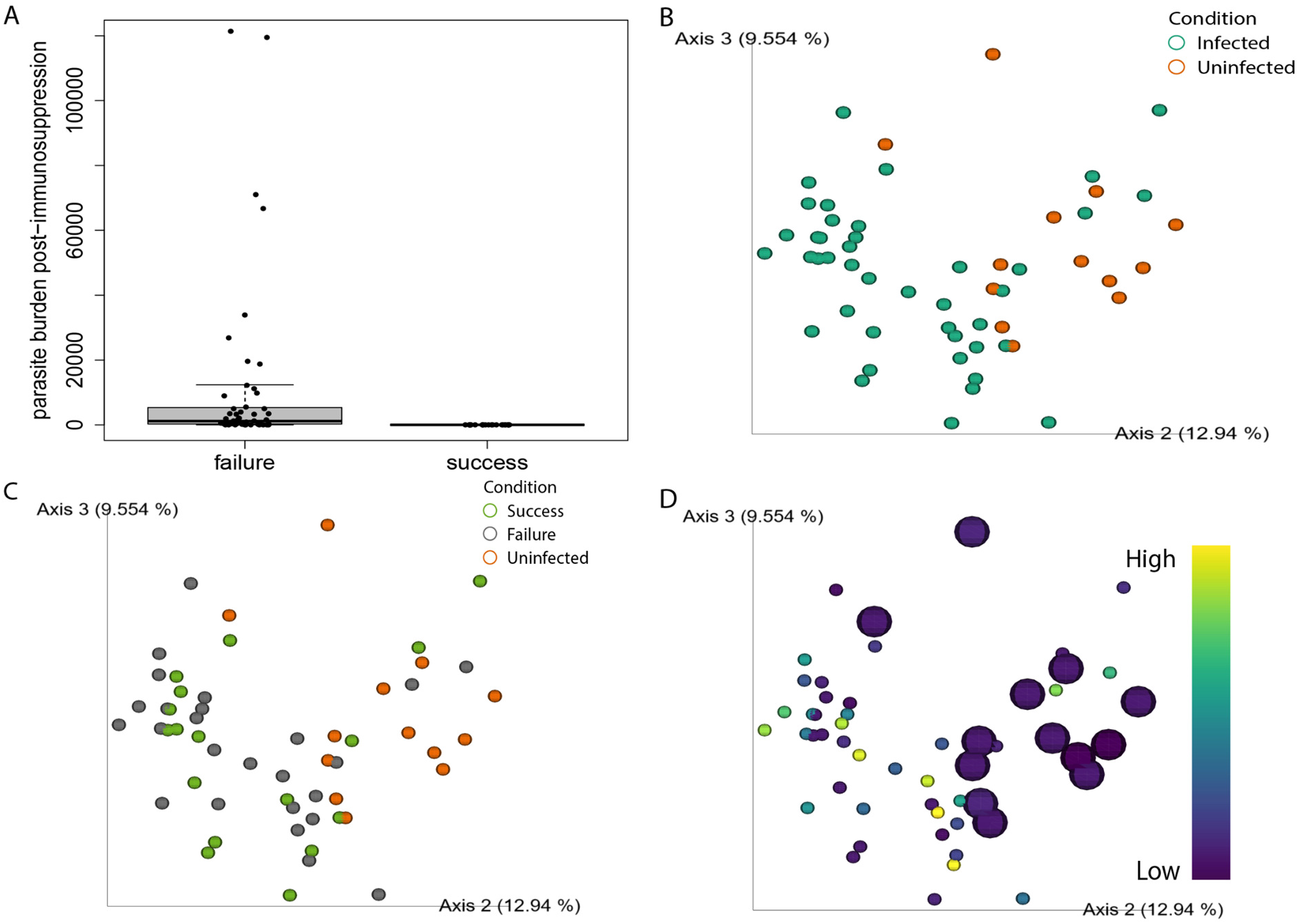
Urine metabolome in mice that successfully cleared *T. cruzi* shows no significant difference from mice in which parasites failed to be cleared. (**A**) Cardiac parasite burden post-infection, treatment and immunosuppression in mice in which parasites persisted (failures) versus those where parasites were successfully cleared (success), as defined by our cutoff compared to qPCR background in uninfected samples. (**B**) Urine sample principal coordinate analysis for mice that were infected and then treated (irrespective of successful or failed parasite clearance) versus uninfected control mice. Samples collected prior to immunosuppression. PERMANOVA p < 0.05 for treated mice versus uninfected mice, R^2^ = 0.08. (**C**) Same PCoA analysis as in (B), recolored to compare successfully-treated mice, mice where treatment failed to clear parasites, and uninfected control mice. PERMANOVA p > 0.05 for successful parasite clearance versus failed parasite clearance, R^2^ = 0.03. (**D**) Same PCoA analysis as in (B), recolored to display normalized cardiac parasite burden post-immunosuppression, R^2^ = 0.02. No segregation by parasite burden was observed. Uninfected controls enlarged for contrast.

**Fig. 6.**
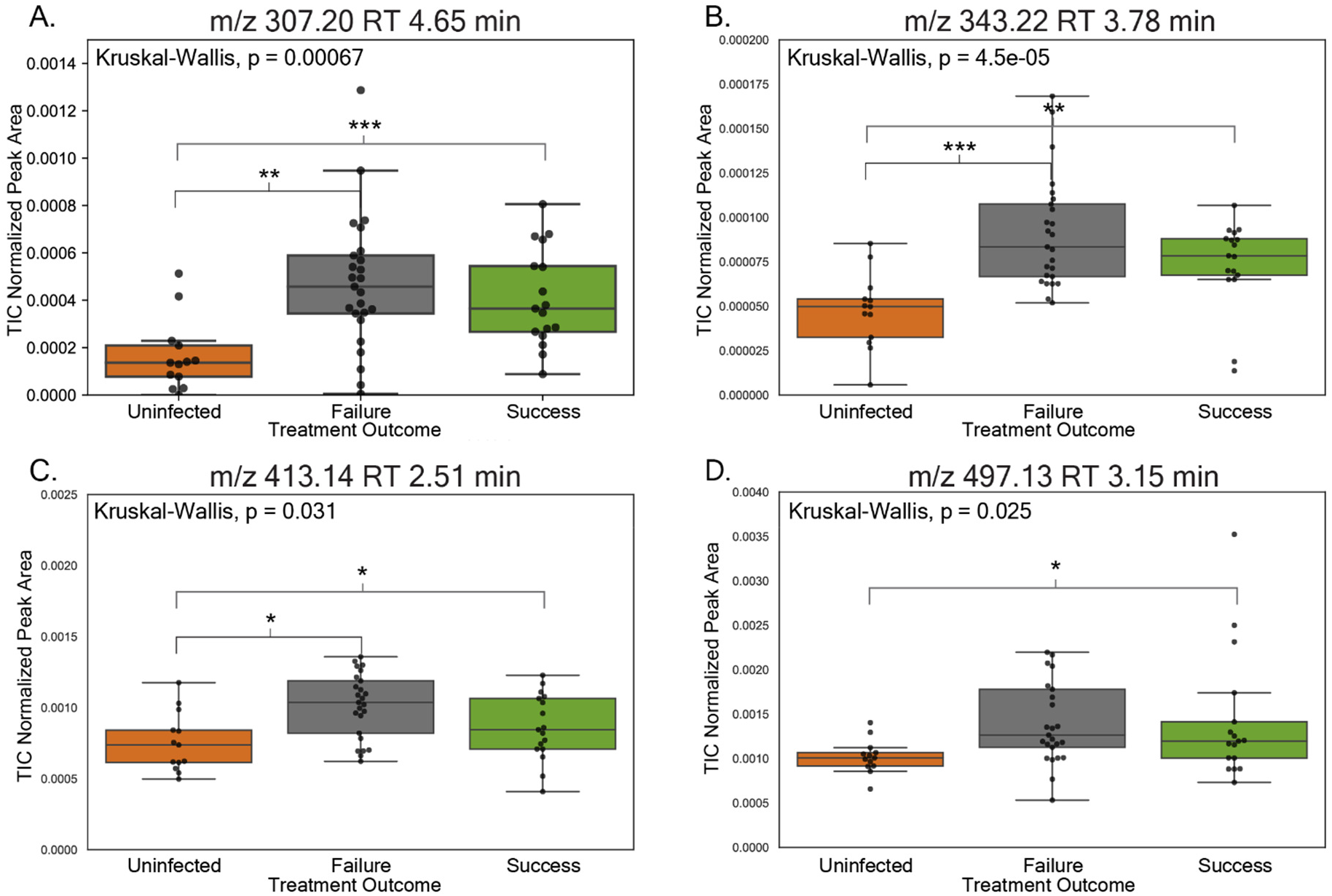
Metabolite features previously identified as differing between infected and uninfected samples remain significantly perturbed even after successful parasite clearance. Significance was established using p-values determined by a Kruskal-Wallis test. P-values were corrected using a Dunn Test with Benjamini-Hochberg adjustment to control the false discovery rate; FDR-corrected p < 0.05 = *, FDR-corrected p < 0.01 = **, FDR-corrected p < 0.001 = ***.

## Discussion

Building on prior work in the heart and in feces ^25, 32^, we analyzed urine, plasma and saliva biofluids with regards to metabolite biomarker discovery for *T. cruzi* infection and antiparasitic treatment success. We found that changes in the urine metabolome were most reflective of disease progression (**Fig. 1**). Furthermore, infected samples diverged from uninfected and isoproterenol-treated samples, indicating that impacts of CD on the urine metabolome diverge from general effects of cardiac damage.

Interestingly, a proof of concept diagnostic test using urine called the Chunap test has been used to determine parasite presence in congenital CD and in *T. cruzi*/HIV co-infected patients ^33, 34^. Both of these studies had increased specificity and sensitivity when compared to current blood-based PCR, microscopy and ELISA methods. *T. cruzi* parasite antigens and DNA have been previously detected in the urine of patients with acute, chronic, and congenital CD ^35–37^. In guinea pigs, it was found that parasite antigens and DNA in urine are not associated with kidney injury or parasites in kidney but rather are an effect of heart and cardiovascular system damage ^38^. Lemos *et al.* identified changes in the kidney in mice infected with high dose (30,000) *T. cruzi* trypomastigotes during the acute stage of the disease ^39^. However, alterations in kidney function were due to cardiac alterations of CD, and the kidneys remained functional ^39^. While kidney damage is seen in CD patients post treatment, there is little research assessing how this damage occurs. Heart failure could be contributing to kidney damage in CD, as is known in other cardiovascular diseases ^40^.These studies signify that our results are not just an artifact of the mouse model and may be well translatable to humans.

Acylcarnitines have already been linked to CD in both heart tissue and the circulation ^25, 41, 42, 43–45^. Our study identified CAR 14:3;O as significantly increased in the urine of males at weeks 4 and 10 post-infection in our validation cohort (**Table 3, Figure S5**). Interestingly, acylcarnitines were decreased by chronic CD in heart tissue in C3H/HeJ mice but increased in acute stage infection in the gastrointestinal tract ^41, 42^. Acylcarnitine metabolism has been causally linked specifically to cardiac metabolism and disease severity in acute *T. cruzi* infection in mouse models ^41, 42^. Lizardo *et al* also identified these classes of molecules as perturbed in the serum by chronic *T. cruzi* infection in mouse models ^45^. Other molecules identified by Lizardo *et al* as potential biomarkers were citrulline, glutamine, spermidine and several sphingolipids ^45^. These metabolites were not identified in our study and may be due to differences in sample type, sample preparation and data acquisition methods, and limits of detection, or had no statistical difference based on time-course analysis.

Based on these results, we sought to use the urinary metabolome to assess treatment success vs failure, using a mouse model with post-treatment immunosuppression as the gold standard of antiparasitic treatment assessment. Interestingly, during chronic CD, while benznidazole treatment did indeed clear parasites in a subset of mice, as expected (**Fig. 5A**), there was no difference in the overall urine metabolome between mice that achieved parasitological cure and those that did not, so that the metabolome of successfully treated infected mice was not reset by antiparasitic treatment (**Fig. 5C**). This contrasts with acute-stage benznidazole treatment, which restored the plasma and heart metabolome ^42^. This could be due to the use of different mouse and parasite strains ^42^, but may also reflect the lack of improvements of clinical outcomes in late-stage patient treatment ^7^ and the disconnect between parasite burden and symptomatic vs asymptomatic chronic infection ^46^. Importantly, our urine results concur with analysis of cardiac tissue in an independent infection model, which likewise showed only minor and incomplete metabolic restoration 56 days post-benznidazole treatment ^47^. As such, our method could represent a quick and non-invasive method to identify compounds with superior efficacy to benznidazole, though this will need to be validated in additional animal models and in human systems. Furthermore, these observations indicate that, while these metabolite alterations are initiated by *T. cruzi* infection, they are markers of CD rather than the parasite itself, and are therefore most likely host-derived. Given that these biomarkers were observed following infection with parasites from two divergent DTUs (TcI and TcVI ^24^) and in two different mouse strains (BALB/c and Swiss Webster), they should have broad applicability. Indeed, when using the Mass Spectrometry Search Tool (MASST) ^48^, 15 out of the 23 (65%) validated biomarkers in Table 3 were present in CD and non-CD human datasets, suggesting applicability beyond mice (**Table S2**).

One limitation is that we only assessed metabolic restoration at 35 days post-treatment. It is possible that metabolism gradually improves over time, though incomplete metabolic restoration was also observed in heart tissue 56 days post-treatment ^47^. A study of CD patients treated with the other antiparasitic nifurtimox showed restoration of many but not all metabolic perturbations up to three years after treatment ^49^. In contrast, the BENEFIT trial that followed patients 7 years post-treatment did not observe any improved clinical outcomes in benznidazole-treated patients ^7^, indicating that longer follow up may not show improvements and that our urine metabolome analysis may be well reflective of clinical outcomes.

## Conclusion

Overall, this study demonstrated a new method to monitor CD progression and to rapidly assess functional treatment success vs failure in chronic CD across multiple DTUs, currently a difficult task. Future work will involve assessment in clinical cohorts. Further investigation of the role of the kidney and of cardiac-renal system crosstalk in CD is also warranted.

## Materials and Methods

### Ethics statement

All vertebrate animal studies were performed under protocol number R17-035 and R20-027, approved by the University of Oklahoma Institutional Animal Care and Use Committee.

### *In vitro* parasite cell culture

*T. cruzi* strain Sylvio X10/4 was obtained from ATCC (catalog number 50800) and placed in co-culture with C2C12 mouse myoblasts (ATCC catalog number CRL-1772). *T. cruzi* luciferase-expressing strain CL Brener was a kind gift of Dr. John Kelly, London School of Hygiene and Tropical Medicine ^30^, and was likewise co-cultured with C2C12 cells. Dulbecco’s Modified Eagle Medium (DMEM) supplemented with 5% iron-supplemented calf serum (Fisher catalog number SH3007203), 100 U/mL penicillin and 100 µg/mL streptomycin was used to maintain cell and parasite cultures at 37°C with 5% CO_2_.

### *In vivo* experimentation – timecourse analysis discovery cohort

5-week-old male Swiss Webster mice (Charles River) were either infected, uninfected, or injected with isoproterenol. Infected mice (n=15) were injected intraperitoneally with 500,000 culture-derived *T. cruzi* trypomastigotes of strain Sylvio X10/4 (TcI DTU ^24^). N=15 mice were left uninfected (mock-injected with DMEM media only). The remaining 15 mice were injected with 100 mg/kg isoproterenol subcutaneously, as a chemically induced heart disease control group_26._

Blood, saliva and urine were collected once a week for 8 weeks and then every other week for 4 weeks. Blood was collected via the saphenous vein and centrifuged for plasma collection. Saliva was collected by intraperitoneal injection of pilocarpine hydrochloride as salivary gland stimulant (0.375 mg/kg) and a cotton swab placed in the mouth for 5 minutes ^50^. The cotton swab was placed into a small eppendorf with the bottom cut and then placed into a large eppendorf. Both eppendorfs with the swab were centrifuged for saliva retrieval (similar to methods optimized in Katemauswa et al ^51^). Urine was collected by placing mice in a single cage until urination. The urine was pipetted into an eppendorf. All samples were stored at –80℃ until extraction.

Infected and uninfected mice (n=3 per group and timepoint) were euthanized at 12 days post infection (acute stage) or 52 days post infection (early chronic stage). All remaining mice from all groups were euthanized at the 90 day endpoint, in the chronic stage of the disease. Isoflurane overdose was used for euthanization. Blood was collected and centrifuged to obtain plasma at all euthanization time points. Phosphate-buffered saline was used to perfuse heart tissue to remove circulating parasites. Hearts were then collected and were transversely sectioned into 4 segments. Heart tissue was stored in RNAlater.

### *In vivo* experimentation – timecourse analysis validation cohort

5-week-old male and female Swiss Webster mice (Charles River) were infected intraperitoneally with 500,000 *T. cruzi* strain Sylvio X10/4 trypomastigotes (TcI DTU ^24^) or remained uninfected (mock-infected with DMEM only; same as for discovery cohort). Urine was collected at week 0, 1 and 2 for both males and females, week 3 for females, week 4 and 5 for males, week 6 for females, week 10 for males and females, week 17 for females, and week 18 for males (**Table 4**). The slight differences in timepoints were for logistical reasons related to other ongoing mouse cohort experimentation.

**Table 4.** Validation cohort mouse numbers per timepoint.

| Week | 0 | 1 | 2 | 3 | 4 | 5 | 6 | 10 | 17 | 18 |
| --- | --- | --- | --- | --- | --- | --- | --- | --- | --- | --- |
| Males, infected | 23 | 24 | 20 | - | 8 | 21 | - | 22 | - | 7 |
| Males, uninfected | 10 | 7 | 8 | - | 5 | 6 | - | 10 | - | 8 |
| Females, infected | 18 | 23 | 23 | 11 | - | 16 | 14 | 15 | 8 | - |
| Females, uninfected | 9 | 10 | 6 | 10 | - | 6 | 3 | 4 | 8 | - |

### *In vivo* experimentation – effects of benznidazole treatment

8-week old female BALB/c mice (Jackson) were obtained and assigned to 2 cohorts: infected (n=45) or uninfected (n=13). Mice in the infected group were injected intraperitoneally with 1,000 luciferase-expressing *T. cruzi* trypomastigotes of strain CL Brener parasites (TcVI DTU ^24^) ^30^, while the uninfected control mice were mock-infected intraperitoneally with DMEM. 116-125 days post-infection (DPI), surviving infected mice were treated with 30 mg/kg benznidazole in solutol once daily via oral gavage. 161 DPI (35 days post-treatment), urine was collected from each surviving mouse in all 3 cohorts using the previously described method. To assess antiparasitic treatment success vs failure, infected mice were then immunosuppressed beginning at 228 DPI (102 days post-treatment) with three rounds of cyclophosphamide, 200 mg/kg by intraperitoneal injection, every 4 days ^52, 53^. 4 days after the final immunosuppression, mice were euthanized and hearts collected to differentiate between successful vs failed parasite clearance by qPCR.

### Parasite burden and cardiac fibrosis quantification

DNA and RNA extraction of heart tissue from the heart base was performed using ZYMO research Quick-DNA/RNA Miniprep Plus Kit (ZD7003), per manufacturer’s protocol. mRNA was reverse transcribed into cDNA using Invitrogen High-Capacity cDNA Reverse Transcription Kit with RNase Inhibitor. cDNA, DNA, and RNA were stored at –20℃ until analysis.

### qPCR

After extraction, DNA was quantified using nanodrop and 180 ng was used for qPCR analysis on a Roche Lightcycler 96 as in McCall *et al* 2017 ^25^, using the conditions in **Table 5** and PowerUp SYBR Green master Mix (Fisher). 2×10^7^ *T. cruzi* trypomastigotes spiked into uninfected samples were used to generate standard curves. Primers used for analysis were ASTCGGCTGATCGTTTTCGA and AATTCCTCCAAGCAGCGGATA for parasite amplification ^54^ and TCCCTCTCATCAGTTCTATGGCCCA and CAGCAAGCATCTATGCACTTAGACCCC for normalization to mouse DNA ^55^.

**Table 5.** qPCR and qRT-PCR parameters.

| Program | Description | Acquisition mode |
| --- | --- | --- |
| Preincubation | 1 cycle |  |
|  | 95°C for 600s | none |
| 3 Step Amplification | 40 cycles |  |
|  | 95°C for 30s | none |
|  | 58°C for 60s | none |
|  | 72°C for 60s | single |
| Melting | 1 cycle |  |
|  | 95°C for 60s | none |
|  | 55°C for 30s | none |
|  | 95°C for 1s | continuous |

### qRT-PCR

qRT-PCR analysis was conducted on a Roche Lightcycler 96. cDNA samples were diluted 1:100 in water and were mixed with PowerUp SYBR Green master Mix (Fisher). Brain natriuretic peptide (Forward: AAGTCCTAGCCAGTCTCCAGA, Reverse: GAGCTGTCTCTGGGCCATTTC) and *Gapdh* primers (Forward: GACTTCAACAGCAACTCCCAC, Reverse: TCCACCACCCTGTTGCTGTA were used for analysis and were added in 0.25 µM concentration ^5657^. qRT-PCR parameters are as in **Table 5**.

### Metabolite extraction and LC-MS/MS data collection (untargeted analysis)

Samples were extracted according to Dunn et al., 2011 ^58^. Briefly, all samples were extracted with 100% HPLC-grade methanol with internal control (0.5 μM sulfachloropyridazine) to a final concentration of 50% methanol (timecourse analysis) or 74% methanol (impact of treatment). Samples were then vortexed and centrifuged at 16,000xg for 15 min. The supernatant was then dried overnight in a Speedvac and stored at –80 °C until LC-MS/MS analysis. For LC-MS/MS analysis, samples were resuspended in 150 μL HPLC-grade water + 0.5 μM sulfadimethoxine, sonicated for 5 min and centrifuged at 16,000xg for 15 min. The supernatant was then transferred to a run plate. Pooled quality controls (QC) were made using 1 μL per well per sample type, and blanks made of resuspension solvent. Samples were analyzed in randomized order per sample type and 10 μL (impact of infection) or 25 μL (impact of treatment) were injected for analysis. LC analysis was performed using a Thermo Scientific Vanquish UHPLC. A Phenomenex 1.6 μm 100 Å Luna Omega Polar C18 column (50 × 2.1 mm) was used for separation. The LC column temperature was set to 40 °C and analysis was done using mobile phase A (water + 0.1% formic acid) and mobile phase B (acetonitrile + 0.1% formic acid) with a 0.5 mL/min flow rate (12.5 min gradient, **Table 6**). Instrumental drift was monitored using a solution of 6 known molecules at the beginning of analysis, between each sample type, and at the end of analysis. Instrument calibration was done using Pierce LTQ Velos ESI positive ion calibration solution immediately prior to instrument analysis. A Q Exactive Plus (Thermo Scientific) high resolution mass spectrometer was used for MS/MS detection (**Table 7**) and ions were generated for MS/MS analysis in positive mode.

**Table 6.** LC gradient.

| Time(min) | Flow (ml/min) | %B | Curve |
| --- | --- | --- | --- |
| 0.000 | Run |  |  |
| 0.000 | 0.500 | 5.0 | 5 |
| 1.000 | 0.500 | 5.0 | 5 |
| 9.000 | 0.500 | 100.0 | 5 |
| 11.000 | 0.500 | 100.0 | 5 |
| 11.500 | 0.500 | 2.0 | 5 |
| 12.500 | 0.500 | 2.0 | 5 |
| 12.500 | Stop Run |  |  |

**Table 7.** Q Exactive Plus (Thermo Scientific) instrument parameters (untargeted data acquisition).

|  |  |
| --- | --- |
| Run time | 0 to 12.5 min |
| Polarity | Positive |
| Default charge state | 1 |
| Full MS |  |
| Resolution | 70,000 |
| AGC target | 1e6 |
| Maximum IT | 246 ms |
| Scan range | 70 to 1050 *m/z* |
| dd-MS2 |  |
| Resolution | 17,500 |
| AGC target | 2e5 |
| Maximum IT | 54 ms |
| Loop count | 5 |
| TopN | 5 |
| Isolation window | 1.0 *m/z* |
| (N)ce/stepped (N)CE | NCE: 20, 40, 60 |
| dd Settings - as in <sup>59</sup> |  |
| Tune Data - as in <sup>59</sup> |  |

### LC-MS/MS data collection (targeted analysis)

An initial untargeted data acquisition run was performed as described above, to obtain total metabolite signal for each sample, and to assess whether any retention time drift occurred compared to the discovery cohort, followed by targeted data acquisition in parallel reaction monitoring (PRM) mode (**Table 8**). Samples were analyzed in randomized order per sample type and 10 μL were injected for analysis. Tune (source) parameters were as in ^59^ and LC parameters were as in **Table 6**.

**Table 8.**
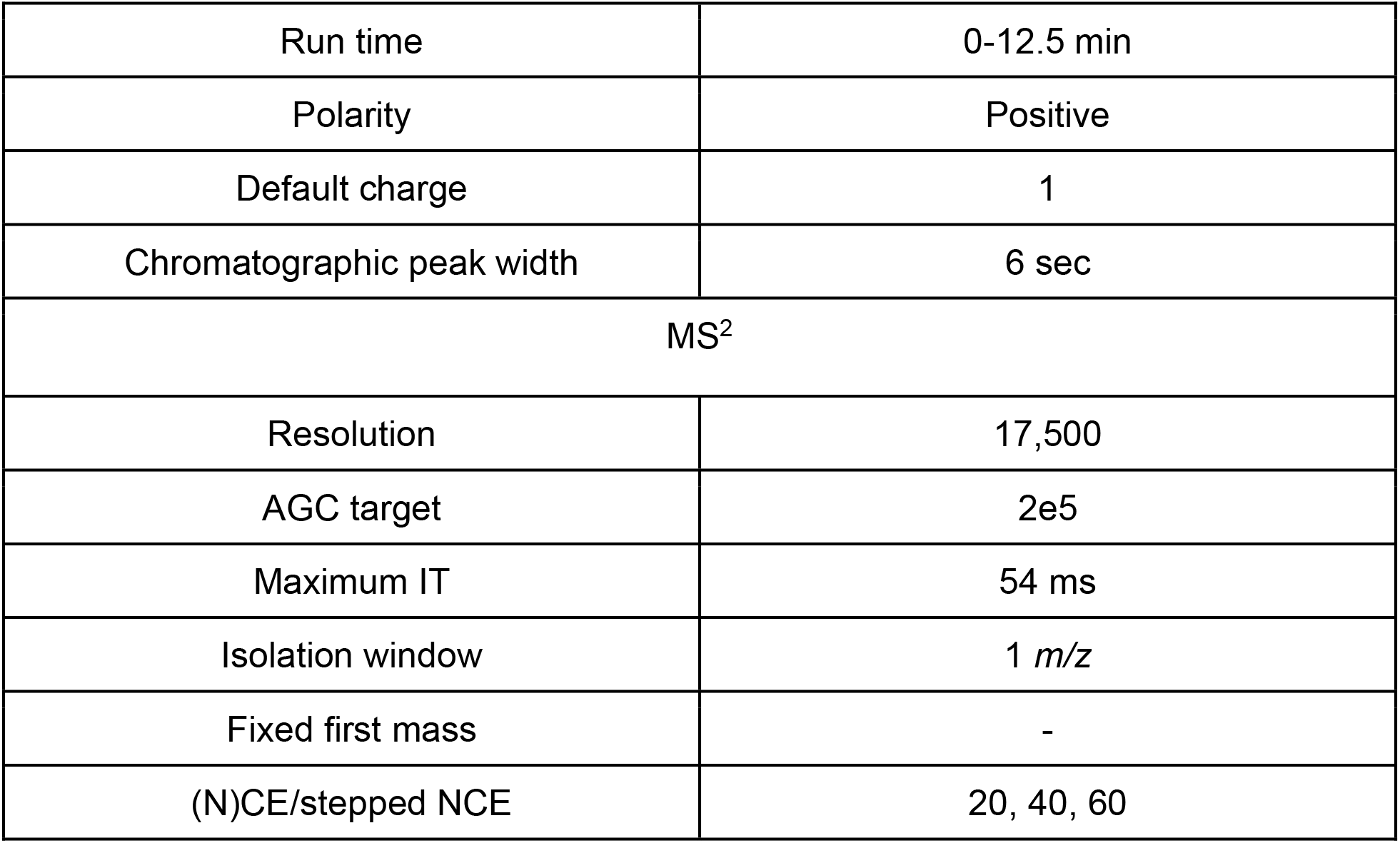
Q Exactive Plus (Thermo Scientific) instrument parameters (targeted data acquisition).

|  |  |
| --- | --- |
| Run time | 0-12.5 min |
| Polarity | Positive |
| Default charge | 1 |
| Chromatographic peak width | 6 sec |
| MS<sup>2</sup> |  |
| Resolution | 17,500 |
| AGC target | 2e5 |
| Maximum IT | 54 ms |
| Isolation window | 1 *m/z* |
| Fixed first mass | - |
| (N)CE/stepped NCE | 20, 40, 60 |

### Data analysis

Data was obtained from LC-MS/MS analysis and converted to mzXML files using MSConvert ^60^. A feature table containing all metabolites for further analysis was developed using MZmine version 2.53 (**Table 9**) ^61^. Three-fold blank removal was performed and total ion current (TIC) normalization performed in Jupyter notebooks in R. We elected not to normalize to creatinine because of evidence that it can be affected by infection status in multiple infection systems (*e.g.* ^62, 6362–6465)^. Principal coordinate analyses (PCoA), longitudinal volatility analyses, and PERMANOVA analyses were all performed in QIIME2 based on TIC normalized features ^66, 67^. PCoA plots were visualized using EMPeror ^68^.

**Table 9.** MZmine 2.53 parameters.

| Type | Parameter | Value |
| --- | --- | --- |
| Feature detection | MS1 Mass Detection | Centroid |
|  | MS1 noise level | 4e5 |
|  | MS2 noise level | 1e3 |
| Chromatogram builder | Masses |  |
|  | Time span | 0.01 min |
|  | Min height | 1.2e6 |
|  | *m/z* tolerance | 10 ppm |
| Chromatogram deconvolution<sup>72</sup> | Local minimum |  |
|  | Chromatographic Threshold | 20% |
|  | Retention Time (minutes) | 0.05<sup>a</sup> ; 1<sup>b</sup> |
|  | Minimum Relative Height | 26%<sup>a</sup> ; 40%<sup>b</sup> |
|  | Minimum Absolute Height | 1.2e6<sup>a</sup> ; 1.0e0<sup>b</sup> |
|  | Minimum Ratio of Peak Top/Edge | 1.19<sup>a</sup> ; 1.20<sup>b</sup> |
|  | Peak Duration Range | 0.01- 1.00 <sup>a</sup> ; 0.04 - 5.00 <sup>b</sup> |
|  | *m/z* MS2 | 0.01 |
|  | RT MS2 (minutes) | 0.1 |
| Isotopic peaks grouper | *m/z* tolerance | 10 ppm |
|  | Retention Time Tolerance (minutes) | 0.1 |
|  | Charge | 3 |
|  | Lowest *m/z* |  |
|  | Monotonic shape | Checked |
| Alignment | Join aligner | 10 ppm |
|  | Retention Time Tolerance (minutes) | 0.3 <sup>a</sup> ; 0.1 <sup>b</sup> |
|  | Weight *m/z* | 1 |
|  | Weight RT | 1 |
| Feature list | Keep only with MS2 scan (discovery cohort only) |  |
|  | Cut off first 20s (discovery cohort only) |  |
|  | Cut off last 30s |  |
|  | Minimum peaks in a row (discovery cohort and treatment effect analysis only) | 8 <sup>a</sup> ; 10 <sup>b</sup> |
|  | Reset ID |  |

<sup>a</sup> Value for timecourse analysis
<sup>b</sup> Value for treatment effect analysis. Where no footnote is provided, values were the same for both analyses.

For comparison of mice with persistent vs cleared parasites, a parasite burden cutoff post-immunosuppression of value in the DMEM group + 3 standard deviations was used (20 parasite equivalents).

Feature-based molecular networking (FBMN) was performed using global natural products social molecular networking (GNPS)^69, 70^ based on the parameters in **Table 10**. Annotations were generated automatically from the GNPS data using an in-house script (https://github.com/camilgosmanov/GNPS ^71^). This provides level 2/3 metabolite annotations based on the Metabolomics Standards Initiative rankings ^29^. The Mass Spectrometry Search Tool (MASST) ^48^ was used to identify the number of human and mouse datasets that contained the validated biomarkers presented in **Table 3**, using the parameters in **Table 11**.

**Table 10.** GNPS Parameters.

| Feature-Based Molecular Network |  |
| --- | --- |
| Precursor Ion Mass Tolerance | 0.02 Da |
| Fragment Ion Mass Tolerance | 0.02 Da |
| Minimum Matched Fragment Ions | 4 |
| Maximum Connected Component Size (Beta) | 100 |
| Maximum shift between precursors | 200 <sup>a</sup> ; 500 <sup>b</sup> |
| Min Pairs Cosine | 0.7 |
| Network TopK | 10 |
| Library Search Minimum Matched Peaks | 4 |
| Search analogs | Do Search |
| Top results to report per query | 1 |
| Score Threshold | 0.7 |
| Maximum analog difference | 200.0 Da <sup>a</sup> ; 100.0 Da <sup>b</sup> |
| Minimum Peak Intensity | 0.0 |
| Filter Precursor Window | Filter |
| Filter peaks in 50 Da Window | Filter |
| Filter Library | Filter Library |
| Normalization per file | Row Sum Normalization (Per file Sum to 1,000,000) |
| Aggregation Method for peak abundances per group | Mean |
| Run Dereplicator | Run |

<sup>a</sup> Value for timecourse analysis
<sup>b</sup> Value for treatment effect analysis. Where no footnote is provided, values were the same for both analyses.

**Table 11.** MASST Parameters.

| Section | Sub-section | Input |
| --- | --- | --- |
| Search Options | Find Related Datasets | Do it |
|  | Parent Mass Tolerance | 0.1 Da |
|  | Minimum Matched Peaks | 4 |
|  | Selected Databases to Search | All |
|  | Ion Tolerance | 0.1 Da |
|  | Score Threshold | 0.7 |
| Advanced Search Options | Library Class | Bronze |
|  | Search UnClustered Data | Don't Search |
|  | Top Hits Per Spectrum | 1 |
|  | Spectral Library | Speclibs |
|  | Search Analogs | Don't Search |
|  | Create Network | No |
| Advanced Filtering Options | Filter StdDev Intensity | 0.0 |
|  | Filter Precursor Window | Filter |
|  | Filter peaks in 50 Da Window | Filter |
|  | Filter SNR Intensity | 0.0 |
|  | Filter Library | Filter Library |

GLMM analysis (Generalized Linear Mixed Model) was performed using the glmmTMB package (Version 1.1.2.3) in the R environment (version 3.6.2) ^73^. Days post-infection and treatment groups (infected, uninfected, isoproterenol treatment) were set as fixed effects while the mouse ID was set as a random effect ^74^. The probability distribution and the link function of the GLMM models were set to Gaussian and identity, respectively. The false discovery rate q-values for the experimental group for each metabolite were derived with the p.adjust function in the R environment.

Targeted data processing was performed using Skyline version 20.1.0.155 ^75^, with parameters similar to those reported in Katemauswa et al., 2022 ^51^ (**Table 12**). Missing values were input as the lowest reported peak area across all samples. Output peak areas were normalized to total sample peak area as determined in MZmine (see **Table 9** for parameters), as an indicator of total metabolite signal. ROC curves were generated in GraphPad Prism 9 on normalized peak areas.

**Table 12.** Skyline parameters.

|  |  |
| --- | --- |
| Prediction |  |
| Precursor mass | Monoisotopic |
| Product ion mass | Monoisotopic |
| Filter |  |
| Precursor adducts | [M+H] |
| Fragment adducts | [M<sup>+</sup>] |
| Ion types | f.p. |
| Auto-select all matching transitions | enabled |
| Library |  |
| Ion match tolerance | 0.5 *m/z* |
| Instrument |  |
| Minimum *m/z* | 50 |
| Maximum *m/z* | 1500 |
| Method match tolerance *m/z* | 0.055 |
| MS1 filtering |  |
| Isotope peaks included | Count |
| Precursor mass analyzer | Orbitrap |
| Peaks | 3 |
| Resolving power | 70,000 at 200 *m/z* |

| MS/MS filtering |  |
| --- | --- |
| Acquisition method | Targeted |
| Product mass analyzer | Orbitrap |
| Resolving power | 17,500 at 200 *m/z* |
| Retention time filtering |  |
| Use only scans within ---- of MS/MS IDs | 5 min |

Fisher’s exact tests were performed using https://www.socscistatistics.com/tests/fisher/default2.aspx. Table of contents graphic created using BioRender.com.

### Data Availability

Metabolomics data has been deposited in MassIVE, accession number MSV000085900 (discovery cohort), MSV000089112 (validation cohort, untargeted), MSV000089113 (validation cohort, targeted), MSV000088436 (effect of chronic-stage treatment). GNPS feature-based molecular networking results can be accessed at https://gnps.ucsd.edu/ProteoSAFe/status.jsp?task=83f9b24689e245f2bf3511cdc29bc45d. GLMM code can be accessed at: https://github.com/thepanlab/timecourseBiomarker.

## Ancillary Information

● *Supporting Information*.

○ Fig. S1. *T. cruzi* infection has minor impacts on the salivary and plasma metabolome.
○ Fig. S2. Representative boxplots of significant metabolites perturbed by infection in the saliva.
○ Fig. S3. Representative boxplots of significant metabolites perturbed by infection or isoproterenol treatment in the plasma.
○ Fig. S4. Representative mirror plots of metabolites perturbed by *T. cruzi* infection or isoproterenol treatment.
○ Fig. S5. TIC-Normalized signal intensity of *m/z* 382.258 RT 4.94 min (annotated as CAR 14:3;O) in male mice across each timepoint in the validation cohort.
○ Table S1. Annotation table of metabolites significantly perturbed by infection or isoproterenol treatment (as identified by GLMM).
○ Table S2. Results of MASST search for the metabolites listed in Table 3.
● *Corresponding Author Information*: Laura-Isobel McCall,
● *Author Contributions*: These authors contributed equally: Danya A. Dean, Jarrod Roach, Rebecca Ulrich vonBargen
● *Acknowledgment*: This project was supported by start-up funds from the University of Oklahoma, with partial support from NIH award number R21AI148886 and R21AI156669, and from the PhRMA foundation, award number 45188. Laura-Isobel McCall, Ph.D. holds an Investigators in the Pathogenesis of Infectious Disease Award from the Burroughs Wellcome Fund. The red-shifted luciferase-expressing *T*. *cruzi* strain CL Brener was a kind gift of J. Kelly, London School of Hygiene & Tropical Medicine, and B. Branchini, Connecticut College. The content is solely the responsibility of the authors and does not necessarily represent the official views of the National Institutes of Health or any of the other funders.
● *Conflicts of Interest:* The authors declare no conflict of interest.
● *Abbreviations Used*:

○ AUC: area under the curve
○ BNP: Brain Natriuretic Peptide
○ CD: Chagas disease
○ DMEM: Dulbecco’s Modified Eagle Medium
○ DPI: days post-infection
○ DTUs: discrete typing units
○ FBMN: feature-based molecular networking
○ FDR: false discovery rate
○ GLMM: Generalized linear mixed models
○ GNPS: global natural product social molecular networking
○ LC-MS/MS: liquid chromatography tandem mass spectrometry
○ *m/z*: mass over charge ratio
○ N/A: not annotated
○ N/S: non-significant
○ PCR: polymerase chain reaction
○ QC: quality control
○ ROC: receiver operating characteristic
○ RT: retention time
○ RT-PCR: real-time polymerase chain reaction
○ TIC: total ion current (normalization)

## Notes

### Competing Interest Statement

The authors have declared no competing interest.

