## Supplemental information for "Persistent biofluid small molecule alterations induced by *Trypanosoma cruzi* infection are not restored by antiparasitic treatment"

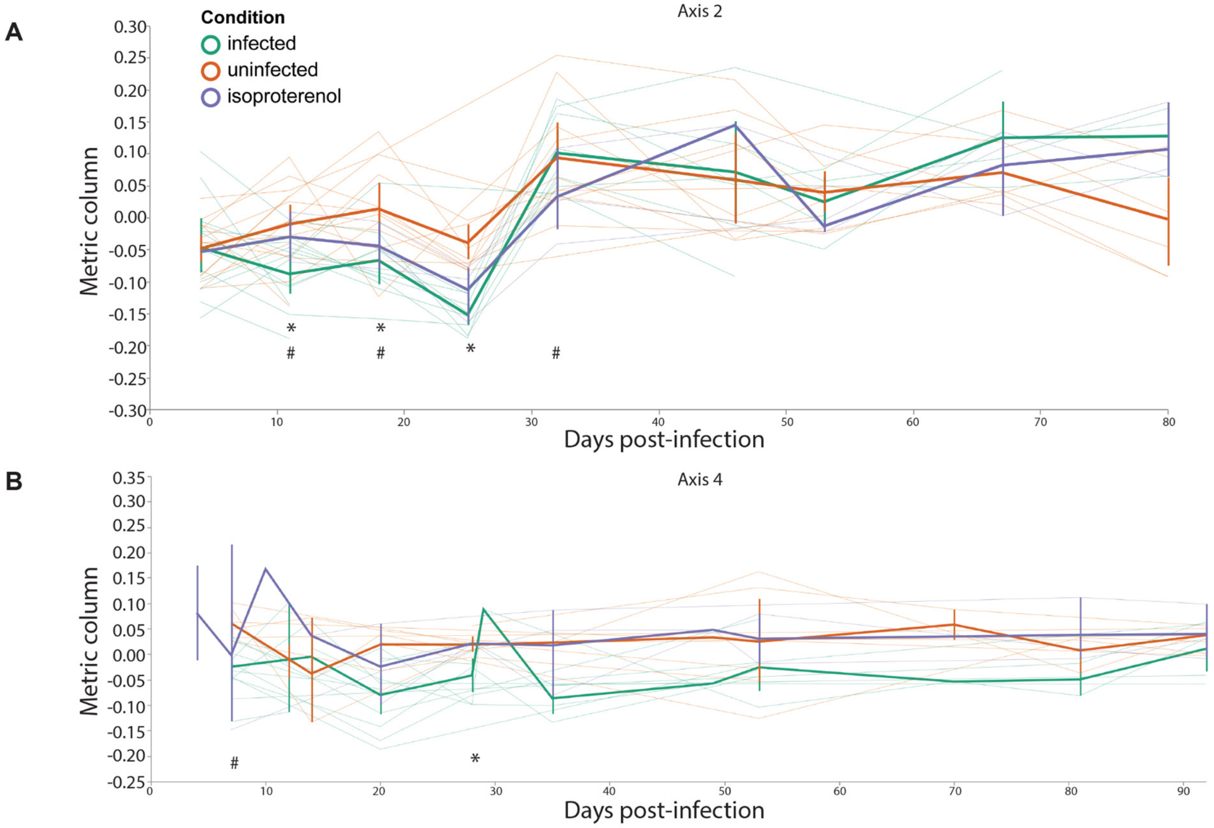


**Fig. S1. *T. cruzi* infection has minor impacts on the salivary and plasma metabolome.** (**A**) Saliva sample volatility analysis showing separation between uninfected samples and both infected and isoproterenol-treated samples at early timepoints along principal coordinate axis 2. *, PERMANOVA $p<0.05$ for infected to uninfected animals. #, PERMANOVA $p<0.05$ for isoproterenol-treated to uninfected animals. (**B**) Plasma sample volatility analysis showing limited differences between groups along principal coordinate axis 4. *, PERMANOVA $p<0.05$ infected vs uninfected. #, PERMANOVA $p<0.05$ for isoproterenol-treated to uninfected animals. Thick lines indicate group mean and thin lines represent the trajectory of each individual mouse along principal coordinate axes.

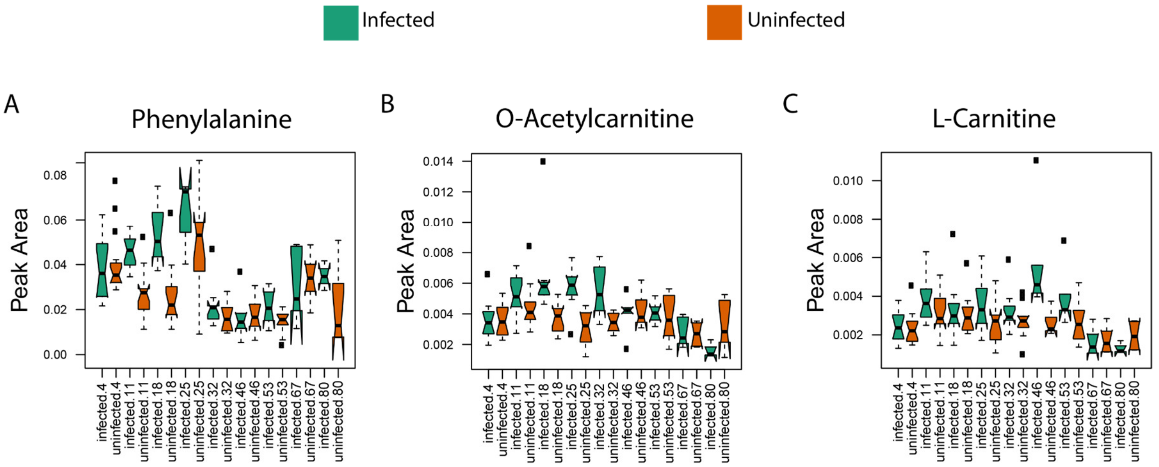


**Fig. S2. Representative boxplots of significant metabolites perturbed by infection in the saliva.** (**A**) Phenylalanine (*m/z* 166.0863, RT 0.63 min). (**B**) O-Acetylcarnitine (*m/z* 204.123 , RT 0.39 min). (**C**) L-Carnitine (*m/z* 162.1124, RT 0.3 min).

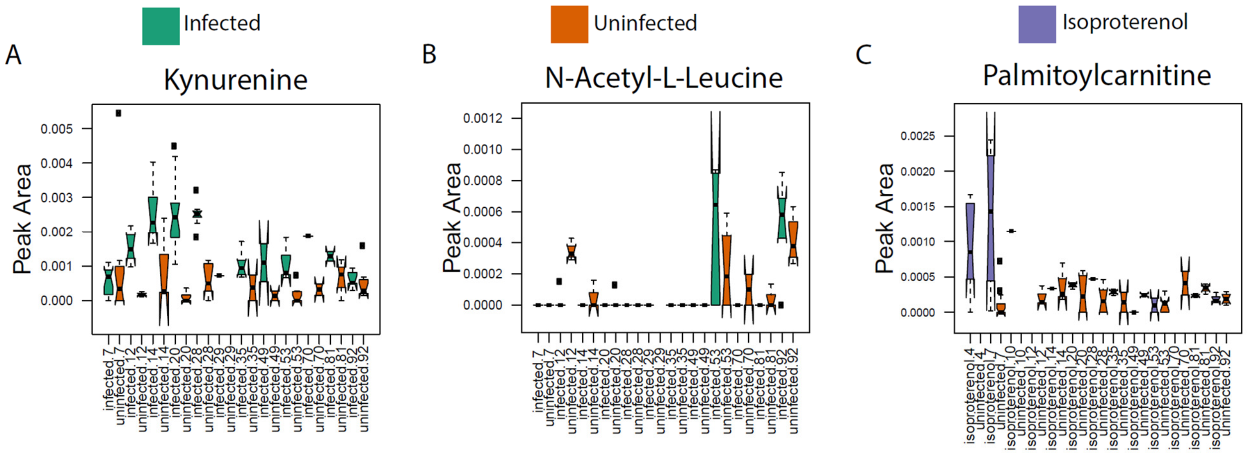


**Fig. S3. Representative boxplots of significant metabolites perturbed by infection or isoproterenol treatment in the plasma.** (**A**) Kynurenine (*m/z* 209.092, RT 0.65 min). (**B**) N-Acetyl-Leucine (*m/z* 174.1125, RT 2.77 min). (**C**) Palmitoylcarnitine(*m/z* 400.3417, RT 6.6 min).

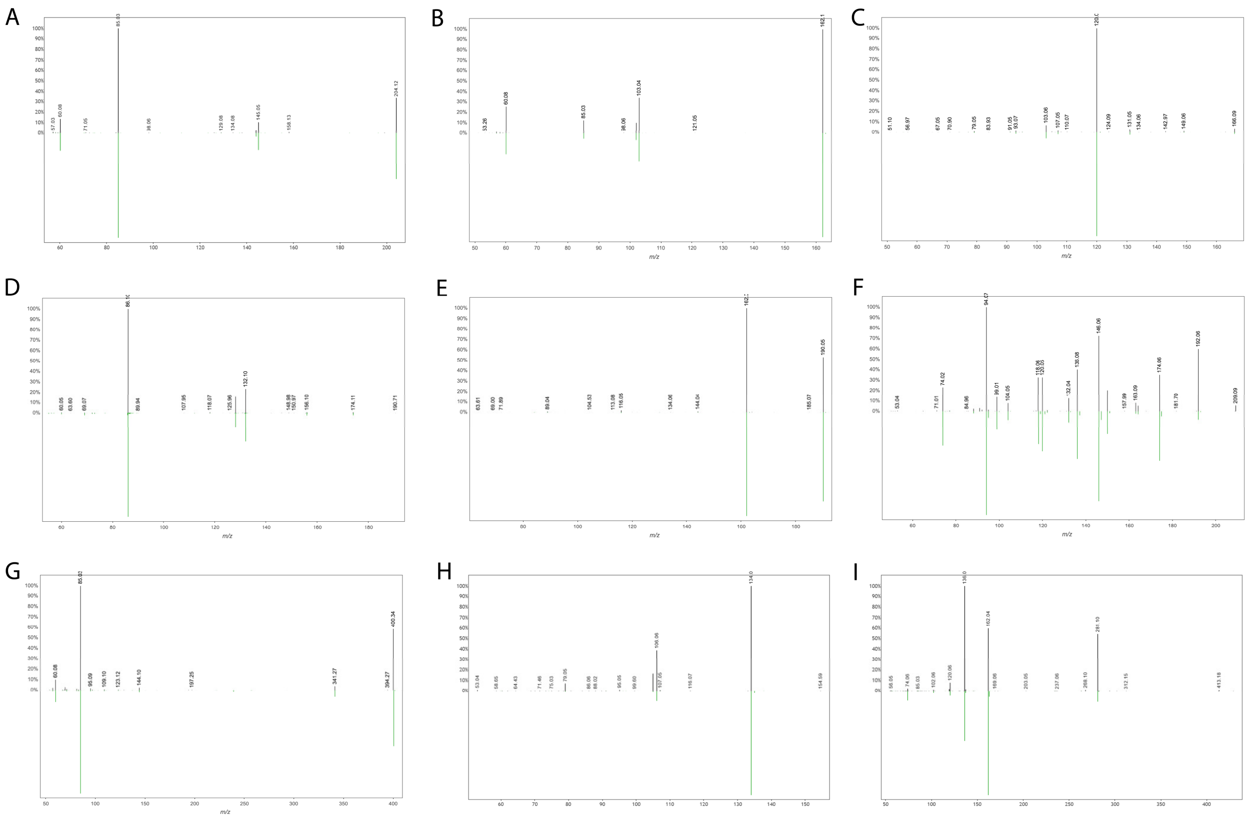


**Fig. S4. Representative mirror plots of metabolites perturbed by *T. cruzi* infection or isoproterenol treatment.** (**A**) Mirror plot of *m/z* 204.123, RT 0.39 min (top, black) to reference library spectrum (acetyl-carnitine, bottom, green). (**B**) Mirror plot of *m/z* 162.1124, RT 0.3 min (top, black) to reference library spectrum (L-carnitine, bottom, green). (**C**) Mirror plot of *m/z* 166.0863, RT 0.63 min (top, black) to reference library spectrum (phenylalanine, bottom, green). (**D**) Mirror plot of *m/z* 174.1125, RT 2.77 min (top, black) to reference library spectrum (N-acetyl-L-leucine, bottom, green). (**E**) Mirror plot of *m/z* 190.0498, RT 2.31 min (top, black) to reference library spectrum (kynurenate, bottom, green). (**F**) Mirror plot of *m/z* 209.092, RT 0.65 min (top, black) to reference library spectrum (kynurenine, bottom, green). (**G**) Mirror plot of *m/z* 400.3417, RT 6.6 min (top, black) to reference library spectrum (palmitoylcarnitine, bottom, green). (**H**) Mirror plot of *m/z* 134.06, RT 1.83 min (top, black) to reference library spectrum (indoxyl sulfate, bottom, green). (**I**) Mirror plot of *m/z* 413.141, RT 2.37 min (top, black) to reference library spectrum (threonylcarbamoyladenosine, bottom, green).

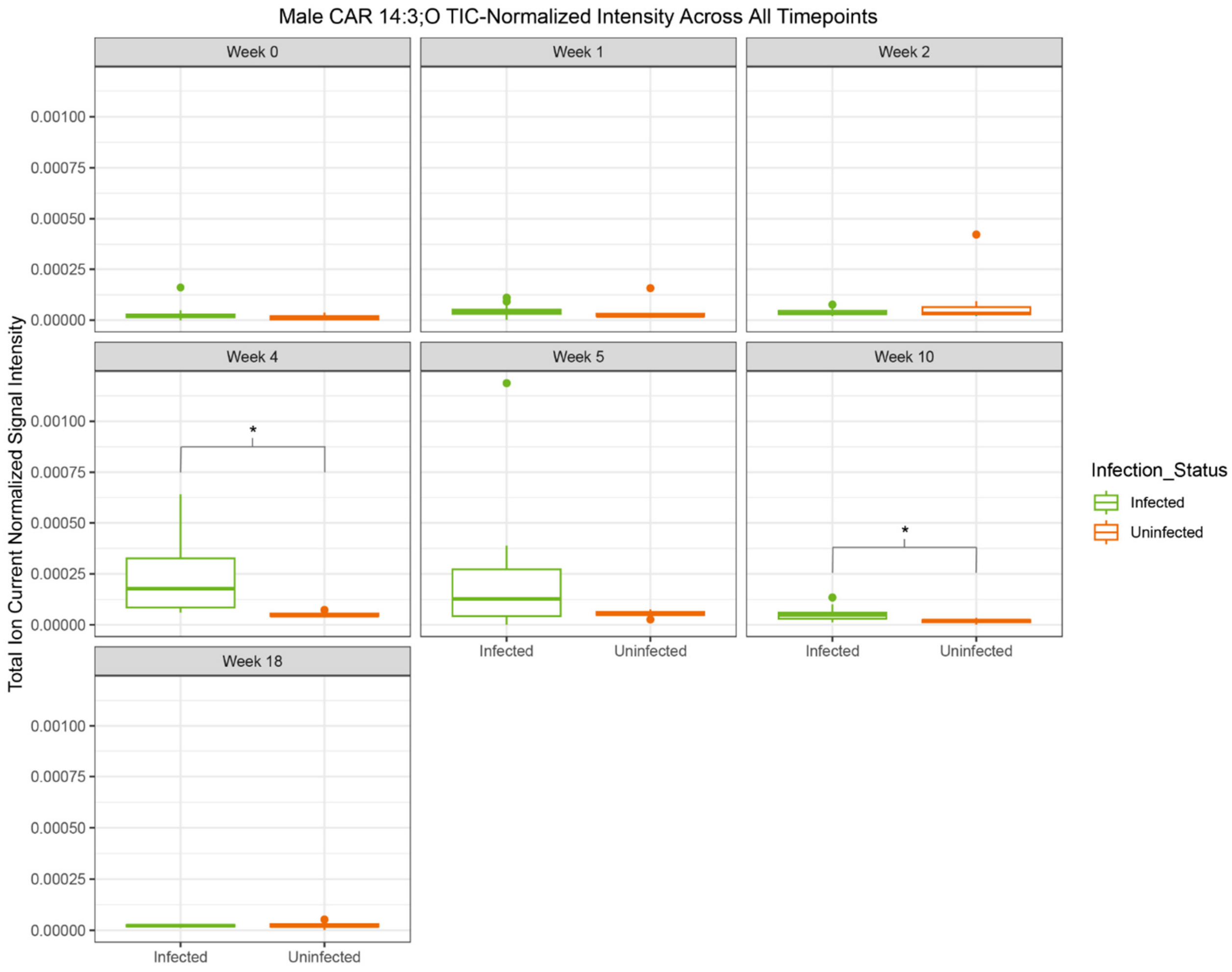


**Fig. S5. TIC-Normalized signal intensity of *m/z* 382.258 RT 4.94 min (annotated as CAR 14:3;O) in male mice across each timepoint in the validation cohort.** *, $p<0.05$ (Mann-Whitney, FDR-corrected).

**Table S1. Annotation table of metabolites significantly perturbed by infection or isoproterenol treatment (as identified by GLMM).**

**Table S2. Results of MASST search for the metabolites listed in Table 3.** Numbers in the "Human" and "Mouse" columns represent the number of datasets that listed the *m/z* of interest as the parent mass and that had MS2 scans that matched our MS2 scans based on 0.7 cosine score threshold. Numbers in the "Total Datasets" column represent the number of all datasets (including human and mouse datasets) that matched the *m/z* of interest.

| *m/z* | Human | Mouse | Total Datasets |
|---|---|---|---|
| 134.06 | 2 | 1 | 4 |
| 151.144 | 1 | 1 | 3 |
| 176.0916 | Not Present | | |
| 214.1071 | Not Present | | |
| 220.0636 | 0 | 5 | 5 |
| 224.1279 | 7 | 1 | 8 |
| 242.1383 | 4 | 2 | 6 |
| 247.1073 | 7 | 8 | 20 |
| 264.0377 | Not Present | | |
| 272.1488 | Not Present | | |
| 280.165 | 0 | 0 | 3 |
| 298.2007 | 7 | 4 | 15 |
| 307.201 | 57 | 27 | 105 |
| 311.069 | 1 | 3 | 5 |
| 333.0509 | 0 | 2 | 3 |
| 343.222 | 6 | 1 | 7 |
| 375.114 | 4 | 1 | 9 |
| 382.258 | 5 | 21 | 33 |
| 400.2687 | 6 | 10 | 17 |
| 411.1543 | 1 | 2 | 3 |
| 413.141 | 53 | 23 | 169 |
| 434.2378 | 0 | 7 | 7 |
| 497.1334 | 1 | 35 | 37 |
